# Replication fork collapse at a protein-DNA roadblock leads to fork reversal, promoted by the RecQ helicase: RecQ promotes replication fork reversal

**DOI:** 10.1101/321869

**Authors:** Georgia M. Weaver, Karla A. Mettrick, Tayla-Ann Corocher, Adam Graham, Ian Grainge

**Affiliations:** School of Environmental and Life Sciences, University of Newcastle, Callaghan, NSW, Australia

**Keywords:** DNA replication, homologous recombination, RecQ, RecG, helicase, replication fork reversal

## Abstract

There are numerous impediments that DNA replication can encounter while copying a genome, including the many proteins that bind DNA. Collapse of the replication fork at a protein roadblock must be dealt with to enable replication to eventually restart; failure to do so efficiently leads to mutation or cell death. Several prospective models have been proposed that process a stalled or collapsed replication fork. This study shows that replication fork reversal (RFR) is the preferred pathway for dealing with a collapsed fork in *Escherichia coli*, along with exonuclease activity that digests the two nascent DNA strands. RFR moves the Y-shaped replication fork DNA away from the site of the blockage and generates a four-way DNA structure, the Holliday junction (HJ). Direct endo-nuclease activity at the replication fork is either slow or does not occur. The protein that had the greatest effect on HJ processing/RFR was found to be the RecQ helicase. RecG and RuvABC both played a lesser role, but did affect the HJ produced: mutations in these known HJ processing enzymes produced longer-lasting HJ intermediates, and delayed replication restart. The SOS response is not induced by the protein-DNA roadblock under these conditions and so does not affect fork processing.

**Author Summary:** To transfer genetic material to progeny, a cell must replicate its DNA accurately and completely. If a cell does not respond appropriately to inhibitors of the DNA replication process, genetic mutation and cell death will occur. Previous works have shown that protein-DNA complexes are the greatest source of replication fork stalling and collapse in bacteria. This work examines how the cell deals with replication fork collapse at a persistent protein blockage, at a specific locus on the chromosome of *Escherichia coli*. Cells were found to process the DNA at the replication fork, moving the branch point away from the site of blockage by replication fork reversal and exonuclease activity. Our data indicate that it is the RecQ helicase that has the main controlling role in this process, and not the proteins RecG and RuvABC, as currently understood. RecQ homologs have been shown to be involved in replication fork processing in eukaryotes and their mutation predisposes humans to genome instability and cancer. Our findings suggest that RecQ proteins could play more important role in replication fork reversal than previously understood, and that this role could be conserved across domains.

## Introduction

During DNA replication, the replication machinery (replisome) can arrest due to impediments on the DNA such as lesions or nucleoprotein blockages. Removal of bound proteins that the replisome itself fails to displace can be carried out by accessory helicases: in E. coli these are Rep, UvrD and/or DinG (1, 2). However, even with the full complement of these helicases, protein roadblocks are still found to be the most common obstacle to replisome progression, especially RNA polymerase (3). Encounter with a protein roadblock can lead to the dissociation of the replisome, and the frequency with which this happens is indicated by the central role of PriA/PriC restart pathways in bacterial cell viability (4). If the blockage is not removed, DNA replication is not able to continue to completion and the cell will not survive. The processing of these stalled forks is likely to be relatively frequent with most or all replisomes predicted to stall during the cell cycle (5, 6). Bacterial replisome dissociation has recently been reported to be occur at a frequency of about 5 events per replisome, per cell cycle (7).

The DNA at a replication fork that has stalled upon encounter with a protein block can be processed by a number of possible pathways. Endonucleases can cut the forked DNA, producing a double strand break, followed by homologous recombination that restores DNA integrity (8). Alternatively, exonucleases can act to degrade the nascent leading and lagging strands, moving the Y-shaped branch point away from the site of blockage (9). Finally, replication fork reversal (RFR) can occur, whereby the leading and lagging nascent strands separate from their respective template strands and anneal to each other, concurrent with the two template strands also reannealing (Reviewed in (10)). This leads to the formation of a four-way DNA structure called a Holliday junction (HJ) that is the substrate for proteins in homologous recombination pathways. This HJ has one arm that has free DNA ends that itself can be acted upon by exonucleases whilst the other 3 arms are continuous DNA.

DNA breakage or the recruitment of RecA to ssDNA at the stalled fork may lead to SOS induction (11). However, the SOS response is only expected to be a major influence in incidences of prolonged DNA damage, high levels of unresolved ssDNA and when the recombination pathways are ineffective at processing blocked forks (reviewed in (12)). RecA acts in the SOS response as a co-protease to cleave the LexA repressor, inducing SOS (reviewed in (13)). The RecA protein has been proposed to play a role in RFR, employing its strand exchange capacity; this is most readily explained as a RecA filament, bound to ssDNA on the lagging strand template of a replication fork, which catalyses invasion into the leading strand duplex, displacing the nascent strand and forming a reversed fork (14). RecA-mediated RFR seems to occur only under specific conditions and is restricted in the presence of the single-stranded DNA binding protein (SSB) (15–17). Other proposed mechanisms that lead to RFR in bacteria, including the involvement of the homologous recombination protein RecA (14), are positive supercoiling ahead of the replication fork (18) and a number of possible helicase/translocase proteins including the branched DNA structure binding proteins, RuvAB and RecG (8, 19–21).

The DNA binding protein RuvA has high specificity towards branched DNA structures and loads the hexameric RuvB onto this DNA to perform the helicase and branch migration activities of the complex (22). When the endonuclease RuvC is also present, the complex is able to resolve the HJ as well as migrate it (23). The monomeric superfamily 2 (SF2) helicase and nucleic acid translocase, RecG, has been shown to be able to migrate HJs but can recognise a wider range of substrates than RuvA. RecG binds dsDNA and has been shown to unwind HJs, D-loops, R-loops and partial fork structures (reviewed by (24)). RecG has a wedge domain that confers its ability to bind to these DNA structures at the ssDNA branch point and two helicase domains that drive the unwinding as the protein translocates (25, 26). The growing evidence for the role of RecG in RFR and the finding that RuvABC-mediated RFR is only required in the absence of certain replisome components has subsequently meant RecG has replaced RuvAB as the major protein implicated in RFR (21, 27, 28). It has, therefore, been suggested RuvAB may only be required to branch migrate the HJ once RecG has performed the initial RFR (21, 29).

RecQ helicases, first identified in E. coli but conserved from bacteria to humans, are monomeric members of the SF2 superfamily (30–32). RecQ can initiate unwinding from a 3’ ssDNA overhang, dsDNA ends, or from internal dsDNA (33–35). RecQ is a multifunctional helicase with roles in homologous recombination and suppressing non-homologous or illegitimate recombination. RecQ has been shown to act with TopoIII as a mechanism for changing catenation of DNA, to counteract illegitimate recombination events and to resolve converging replication forks (36, 37). During homologous recombination the helicase is predominantly associated with the RecFOR recombination pathway employed to process ssDNA gaps in the DNA commonly caused by UV irradiation (38, 39). In this pathway, RecQ translocates along the ssDNA gap and unwinds dsDNA, further enlarging the single stranded region so that other proteins can gain access to the DNA damage, including the RecFOR proteins that can then load RecA. In vitro analysis of the activity of the human RecQ helicases BLM and

WRN on model fork structures determined that they are both able to reverse forks into HJs (40, 41). However, *E. coli* RecQ is yet to be implicated in RFR *in vivo*. Interestingly, BLM and WRN can also promote the opposite reaction, un-reversing HJs back into forks (42). Another human RecQ homologue, RECQ1, has been shown to be important in vivo for the restart of replication forks that have been reversed and catalyses restoration of forks from their reversed state (43).

One of the major impediments to studying RFR in vivo has been the combination of a large chromosome and a rapid replisome: E. coli contains a genome of ∼4.6 Mbp and the replisome moves at a rate close to 1kb/s (4). A system has been developed whereby a site-specific road block to DNA replication can be induced at a known position in the chromosome using a transcriptional repressor (TetR) bound to an array of operator sites within the *E. coli* chromosome (44, 45). Addition of a temperature sensitive allele of the replicative helicase (DnaBts) allows the rapid inactivation of the replisome across a population of cells, and the replication forks are processed by the cell within 5 minutes (44). Furthermore, the disappearance of the forked DNA structure from the array region coincided with the visualisation of HJs upstream of the array suggesting RFR had taken place in a sizeable proportion of cells (44). The disappearance of Y-shaped DNA will be used here as a definition of replication fork collapse; whether or not this is accompanied by partial or complete dissociation of the replisome is unknown. The regression of the replication fork away from the site of DNA damage or protein block allows repair proteins or accessory helicases to access the DNA and resolve the problem. The upstream HJ that is formed may be processed in a recombination-dependent or independent manner to restore a replication fork (reviewed by (10)). If the blockage is removed, replication is able to restart using the reformed fork structure onto which replisome reloading takes place (45), most likely in a PriA-dependent manner. Indeed, replication was observed to restart and proceed through the array in the vast majority of cells within 5 minutes of the addition of release of the block by addition of anhydrotetracycline.

In this report a site-specific replication block was established and characterised at the lac locus, approximately midway round the right replichore on the *E. coli* chromosome. A *dnaBts* allele was introduced to be able to trigger replication fork collapse by shifting to a non-permissive temperature. This site-specific replication blockage system was used to examine the process of RFR, and the relative contributions of candidate proteins. RFR is seen to occur as a major pathway used in cells to deal with a persistent roadblock to replication, with little or no evidence for endonuclease action at the fork. Some exonuclease activity is seen as well, that could either be from action directly at the Y-shaped fork, or at the HJ produced by RFR. Deletion of recQ is seen to have the greatest effect on RFR with the process being highly inefficient in its absence, while both ruvABC and *recG play* a more minor role in RFR. However, the data suggest that RecG is involved in migrating the HJ formed by RFR back into a forked DNA structure that can be used for replication restart; in the absence of RecG HJs are much more prominent and are seen to persist. The induction of the SOS response was only observed to begin after 4 hours of replication fork blockage, implying that it does not affect fork processing during the standard experimental timeframe used here.

## Results

### Visualising replication fork reversal by inducing replisome collapse

Previous work has established that a fluorescent repressor/operator system (FROS) using an array of *tetO* sequences integrated 16kb anticlockwise from oriC can block replication when the cognate repressor (TetR-YFP) is overproduced (44, 45). In this study a similar tetO array is utilised, inserted at the lac region roughly half-way between the origin and the terminus on the clockwise replichore. *To verify that the roadblock was functional at* the new position a set of growth experiments were carried out to determine whether replication could be blocked efficiently across the population, and whether this blockage was fully reversible (Fig. 1A). The induction of TetR-YFP with arabinose for 2 hours at 30°C resulted in the vast majority of cells (98%) containing one fluorescent focus per cell, determined microscopically (Fig. 1A). Cells with a single focus, representing the tetO array, have not duplicated the array region. When replication is not blocked in the unsynchronised culture, there is a mixed population of cells with one, two and sometimes more than two foci. A predominance of cells with a single focus point indicates replication has been arrested across the population, and is used in subsequent experiments as an indication that replication blocking has been successfully established prior to further experimentation (44, 45). The viability of these cells following blockage of replication was examined by plating cultures on LB agar and the colony forming units were determined. Cells that had been grown with arabinose to induce the DNA replication roadblock had ∼1000-fold decrease in viability compared to untreated cells (Fig. 1B).

**Fig 1:**
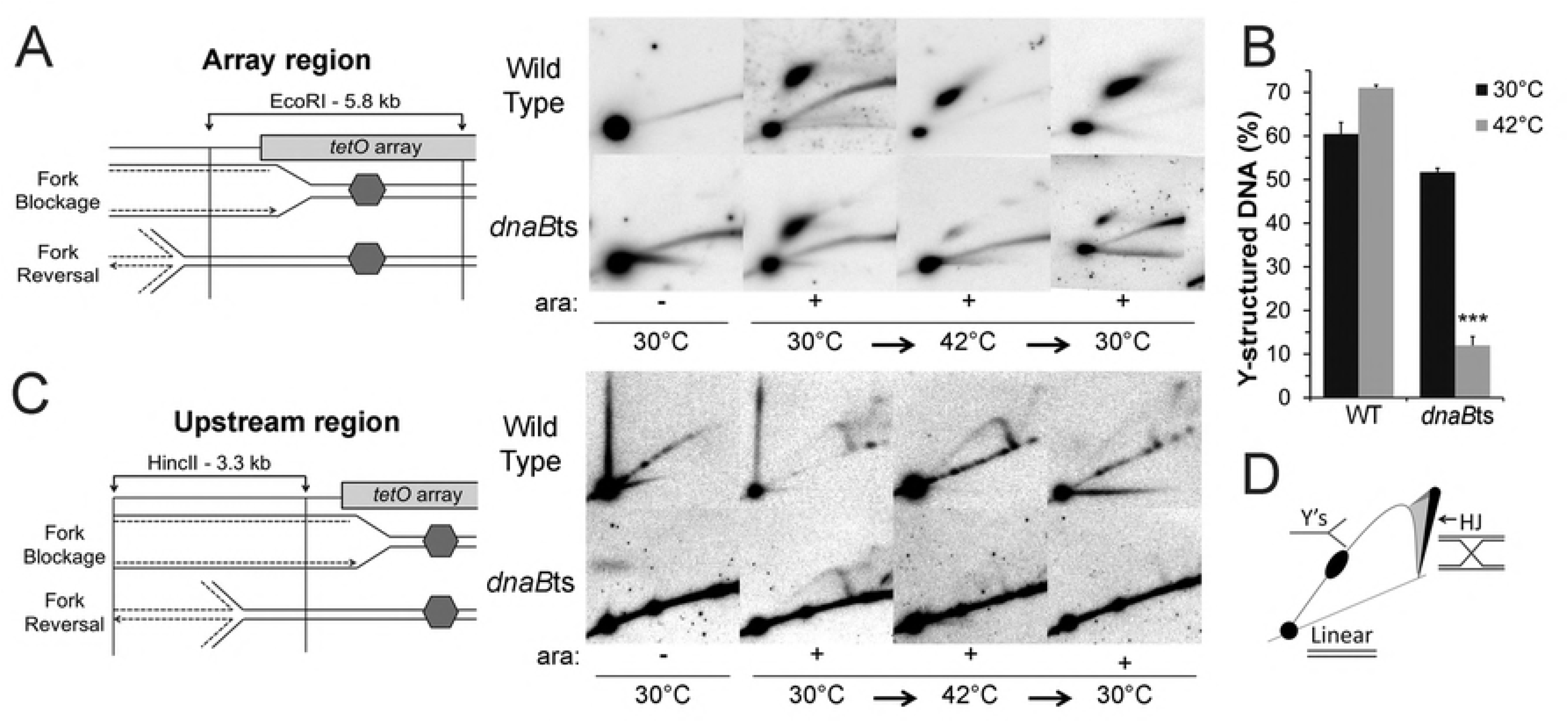
TetR binding to a tetO array creates a controlled replication block that is also reversible. Wild type and a strain carrying dnaBts were grown at 30°C with 0.1% arabinose (ara) for 2 hours and a subset of these cells were treated with anhydrotetracycline (AT) for 10 minutes. The arabinose treated cells were shifted to 42°C for an hour (the non-permissive temperature for DnaBts) then returned to 30°C for 10 minutes (TS). A subpopulation of these cells were treated with AT. Percentage of (*A*) Wild type or (*C)* DnaBts cells containing one focus or multiple foci after specified treatments, formed by the overproduction of TetR-YFP. Viability of (*B*) Wild Type or (*D*) DnaBts following indicated treatments. A 10-fold serial dilution of the cells was conducted and spotted or spread onto agar plates supplemented with Amp (for arabinose treated cells), or Amp and AT (for AT treated cells). DnaBts cells were grown on agar plates that also contained Kan. Spot plates: colony growth at indicated dilutions. Graphs: cell viability shown as colony forming units per mL +/- SEM. Number of biological replicates: 4.

To assess whether the established replication block could be removed to allow the resumption of replication, anhydrotetracycline (AT) was added to blocked cells and cultures were reanalysed after a 10 minute period. 80% of the cells now contained two or more fluorescent foci, indicating that the array had been duplicated and that the two copies had segregated from each other (Fig. 1A). Given the position of the array, the 10 minute AT exposure does not allow time for a newly formed fork to start from oriC and progress to the array position. Thus, the multiple foci are most likely produced exclusively from restarted forks. The viability of these cells was restored to initial untreated levels, confirming that replication was able to restart throughout the population (Fig. 1B). These results confirm that the *tetO* array, incorporated at *lac*, is an effective blockage to replication when the repressor is overproduced, and that the roadblock is reversible with addition of AT.

*dnaBts* was incorporated into the *tetO* array-carrying strain and assessed in comparison to a WT strain for its ability to block and restart replication (Fig. 1C, 1D). It has recently been proposed that the replication fork present at the TetR-YFP roadblock is not stable but has a half-life of 3–5 minutes (44). An equilibrium exists between forks that are stalled at the block and ones that have collapsed and are being processed and will subsequently restart leading to their collision with the replication roadblock once again. DnaBts is utilised here to ensure the synchronous dissociation of the replisome across the entire population and to provide a regulatory tool where replication restart is dependent on DnaBts reactivation. The incorporation of dnaBts did not alter the ability of the replication roadblock to function at permissive temperature, and the results were identical to the WT strain (Compare Fig. 1A to Fig. 1C); following overproduction of the TetR-YFP repressor 98% of cells had a single fluorescent focus and viability was decreased over 1000 fold. Addition of AT, to release the replication roadblock, led to the rapid restart of replication with 80% of cells containing 2 or more foci after 10 minutes, and a complete recovery of viability.

Following validation of the replication block in both strains, the effect of a shift to 42°C, a non-permissive temperature for DnaBts, was investigated. Cells were grown at 30°C and replication was blocked by overproduction of TetR-YFP as before. Each strain was then transferred to 42°C for one hour, to inactivate DnaB in the temperature sensitive strain. The ability of DNA replication to restart in each strain was then compared by transferring cells back to the permissive temperature of 30°C for 10 minutes, with or without the addition of AT. In the WT strain a similar pattern was seen to cells grown only at 30°C; the majority of the population had a single fluorescent focus following the temperature shifts indicating replication was still being efficiently blocked, and addition of AT led to ∼70% of cells showing two or more foci within 10 minutes of addition of AT demonstrating effective replication restart (Fig. 1A). Similarly, viability was almost fully recovered by addition of AT. Therefore, the temperature shift to 42°C had little noticeable effect upon the replication block or replication restart in the WT strain. In the *dnaBts* strain the 1 hour at 42°C did not affect the integrity of the replication blockage as judged by the high proportion of cells with a single focus and the drop in viability when arabinose is present (Fig. 1C, 1D). Upon addition of AT when the cells were returned to 30°C viability was also seen to recover to WT levels. This demonstrates that transient inactivation of DnaBts is well tolerated and cells recover fully when returned to a permissive temperature and the replication blockage is removed. However, the proportion of cells with 2 or more foci after 10 minutes of AT treatment is much lower than seen with WT. This likely reflects the extra time this strain requires to restart replication that may involve the refolding of inactive DnaBts protein or the novel synthesis of the protein. It has been shown that the majority of locations on the E. coli chromosome, including lac, require 7–10 minutes to visibly segregate from the sister DNA following replication (46, 47). Thus, even a short delay in restarting replication in the dnaBts strain may be sufficient to prevent the replicated sister duplexes from separating from each other, within the resolution limit of the microscope.

The DNA structures present within each strain during the replication block experiment were visualised using 2-D neutral-neutral agarose gels (Fig. 2). DNA from each condition was extracted, digested and electrophoresed. A radiolabelled probe was used to detect either a 5.8 kb array region including 0.6 kb upstream from the beginning of the array, or a 3.3 kb region from 0.5 kb to 3.8 kb upstream of the array (Fig. 2). At 30°C, both the WT and dnaBts strains showed a strong, localised spot of Y-shaped DNA in the array region after the addition of arabinose, indicating replication fork blockage (Fig. 2A). With a shift to 42°C for an hour, persistent forked-DNA structures remained in the WT strain consistent with maintaining the equilibrium of blocked forks and fork turnover (Fig. 2A) (44). The additional hour of blockage and the temperature shift did not significantly alter the replication blockage in the array region of the WT strain. Quantification of the linear and forked DNA signals determined that ∼60% of DNA was seen to be Y-shaped at 30°C; ∼70% was as a Y-structure after the hour at 42°C (Fig. 2B). The region upstream of the array showed that upon replication blockage a clear signal for HJ and Ys could be seen (Fig. 2C), and the levels of these signals did not vary with changes in temperature in the WT strain, suggesting that the replication forks maintain a relatively constant fork turnover rate.

**Fig 2:**
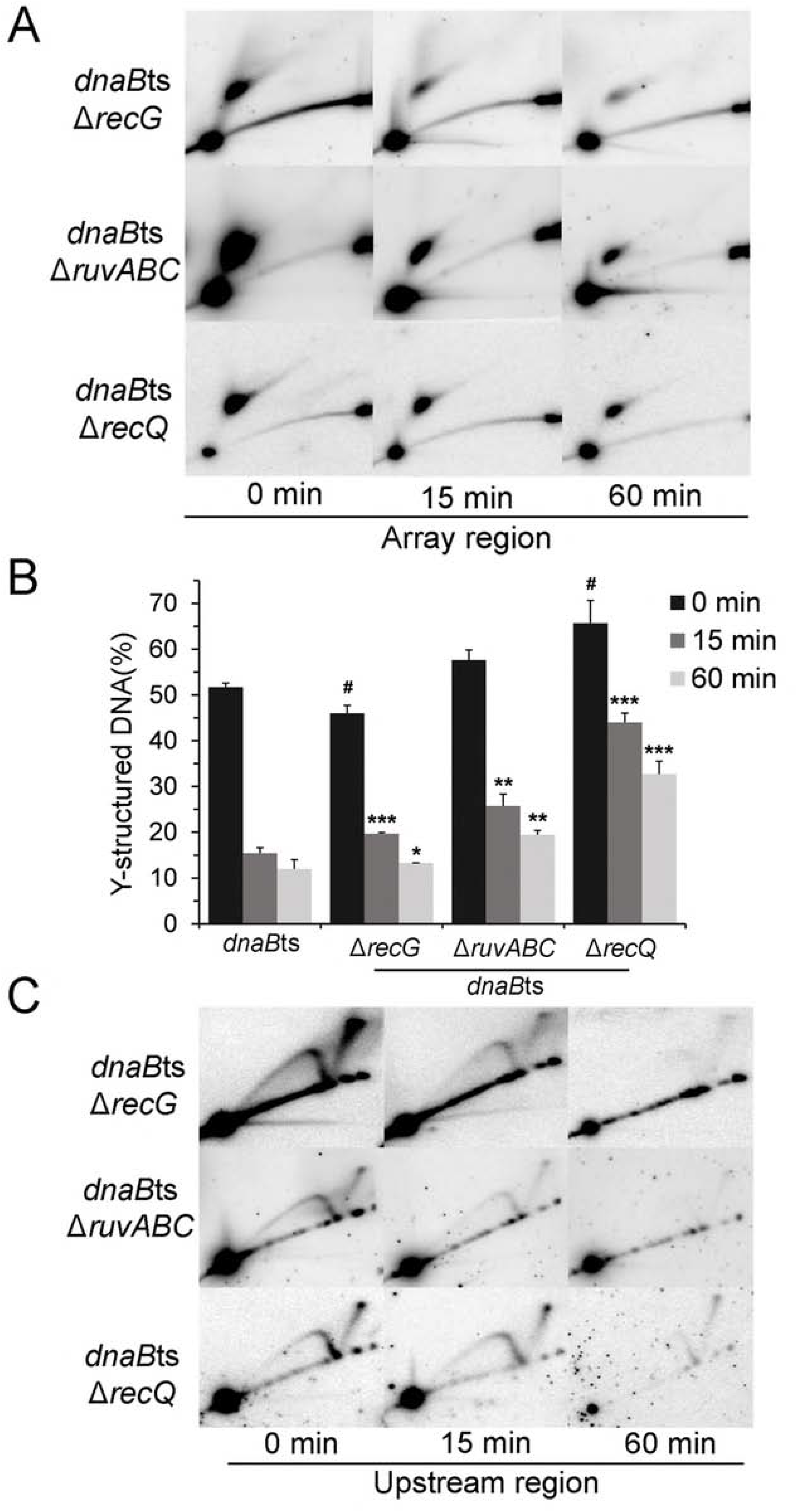
Replication fork collapse visualised by 2-D agarose gel electrophoresis. (A) DNA structures from indicated conditions for array region in WT and DnaBts strains, visualised by Southern hybridisation. Schematic represents array fragment created by digestion with EcoRI (arrows indicate restriction sites). (*B*) Quantification of the Y-structured DNA in the array region at 30°C (2 hours growth with arabinose) and 42°C (an additional 1 hour grown with ara). *** = significantly different (*P* < 0.001) from its 30°C counterpart (*C)* DNA structures from indicated conditions for region upstream of array in WT and DnaBts strains, visualised by Southern hybridisation. Schematic indicates upstream fragment created by digestion with HincII (arrows indicate restriction sites). (*D*) Representative image illustrating the various DNA structures detected by the probes. When replication is blocked, forks remain in the array fragment and accumulate at a similar position on the Y-arc. A line signal protruding from the linear DNA and/or a cone shape extending from the very top of the Y-arc represent Holliday junctions (HJ). See also Fig. S1. n = 2–3

In contrast, the *dnaBts* strain displayed a vast decrease from ∼50% to 12% in forked DNA signal at the array following the shift to 42°C (Fig. 2A and B). The non-permissive temperature for DnaBts induced the collapse of the replication fork and resulted in fork processing events, including RFR, moving the forks out of the array region, as visualised by the dramatic loss of Y-structured DNA. Upstream of the array HJ and Y-shaped signals are seen at 30°C as with WT (Fig. 2C). After 1 hour at 42°C mostly linear DNA remained upstream suggesting that the reversed forks had been further processed, either by migration further upstream or by nuclease action (Fig. 2C; Fig. S1); replisome reloading cannot occur in the dnaBts strain at 42°C explaining the difference compared to WT.

When the *dnaBts* cells were returned to the permissive temperature of 30°C for 10 minutes the forked DNA signal at the array increased above the level seen at 42°C, consistent with restoration of DnaB function, restart of replication and subsequent collision with the block (Fig. 2A). However, the amount of Y-shaped DNA was lower than the level seen prior to the shift to 42°C, possibly due to the time taken to reactivate/re-synthesise DnaB and then to restart the replication fork. This is in agreement with the microscopy data shown in Fig. 1.

### Induction of the SOS response by persistently blocked replication forks begins after 4 hours

It was possible that the SOS response may have been activated in these cells after prolonged replication fork stalling events (∼2 hours), as used in this study. In order to address this concern, the timing and extent of SOS response was monitored in cells that had replication blocked (and released). sulA is a mid-stage gene in the SOS regulatory network with its activation occurring 5 minutes after UV irradiation, with a 100-fold increase in transcription once SOS is induced (48, 49). The *sulA* promoter was cloned to drive expression of the mCherry fluorescent protein, and this construct was inserted half way round the left replichore on the chromosome. mCherry fluorescence would be indicative of the *sulA* promoter activation and, therefore, SOS induction. The fluorophore fused to TetR in this strain was also changed from YFP to GFP to reduce spectral emission overlap with mCherry. mCherry fluorescence in cells was measured after 0, 2, 4, 6 and 24 hours of arabinose induction of the TetR-GFP roadblock using a microplate reader and visualised by fluorescence microscopy (Figure 3) to determine whether the SOS response is induced by a persistent replication fork blockage. This reporter psulAmCherry construct was also incorporated into both *recA* or *uvrD* strains as controls; in *recA* cells the SOS response cannot be induced, while in a *uvrD* mutant the SOS response cannot be repressed once induced.

**Fig 3:**
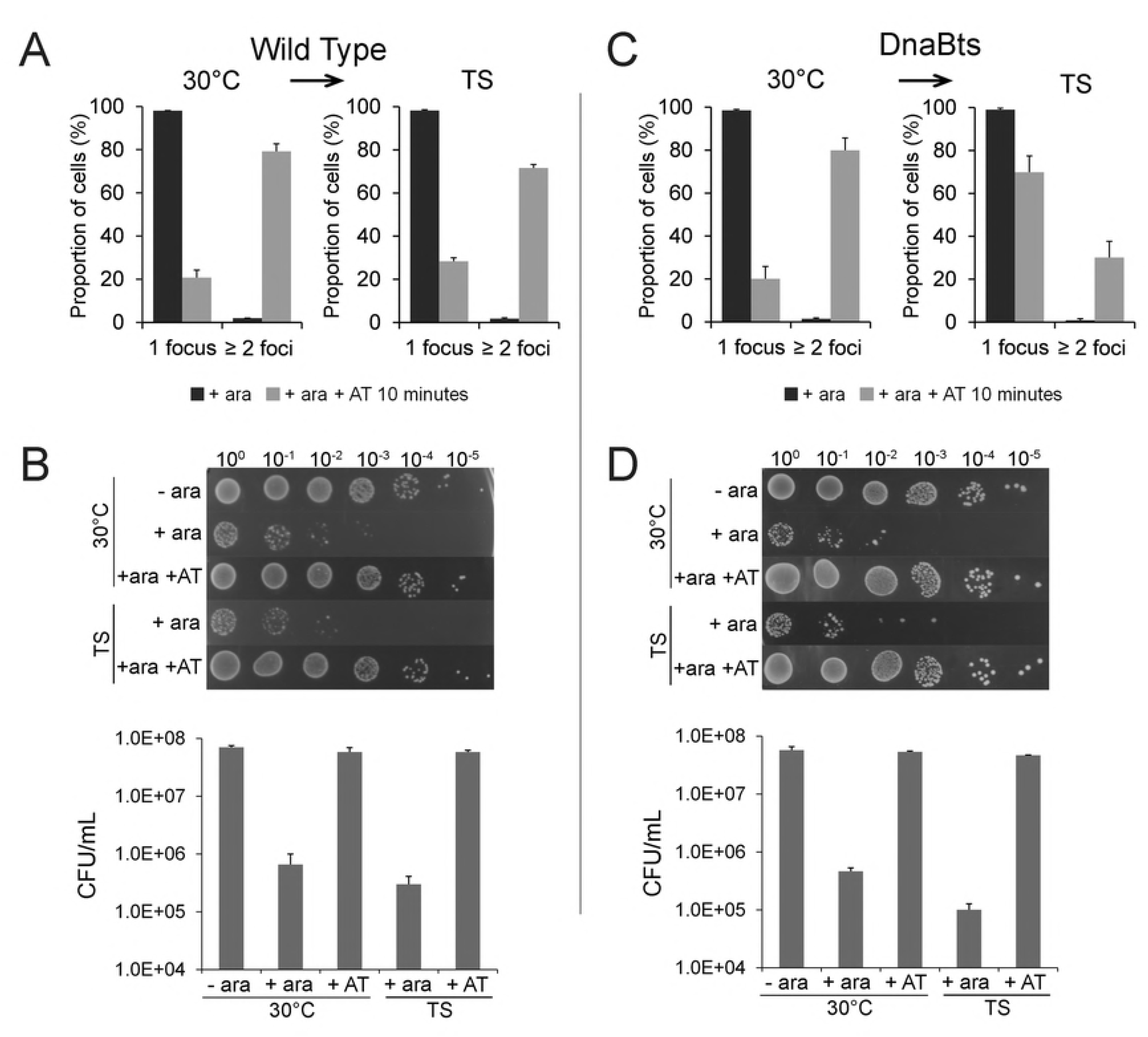
SOS induction begins after 4 hours of a persistent replication block and increases thereafter. The strain carrying sulAp-mCherry was grown with 0.05% arabinose at 30°C and samples were taken after 2, 4, 6 and 24 hours. After the 2 hours, a subset of cells were incubated with AT (10 µg/mL) and samples taken at 2, 4 and 22 hours. SOS induction was observed via A) mCherry fluorescence emission at indicated time points in Relative Fluorescence Units (RFUs) in blocked (+ara) and released (+ara +AT) cells. Average and standard deviation is shown from 3 biological replicates. B) microscope images at each time-point in replication blocked cells showing an overlay of phase contrast images and mCherry fluorescence. Red cells indicate *sulAp*-mCherry expression (SOS induction) Scale bar = 4 microns. See also Fig. S2.

SOS induction had not occurred 2 hours after the induction of the replication fork block in WT cells, as mCherry fluorescence was not detected by either microplate reader or microscopy (Figure 3A and B). However, blocked replication forks had been established across the cell population as indicated by the single GFP focus points in each cell (Fig. S2). Some evidence of SOS induction was observed at 2 hours in uvrD cells though not in recA (Fig. S2). 4 hours after induction of the replication fork block, each strain still had a single focus across the population confirming replication could proceed through the array, and cells had begun to elongate (Fig. 3 and Fig. S2). At this time point, some cells showed delocalised mCherry fluorescence by microscopy, and the population showed a slight increase in red fluorescence detected by the plate-reader (2250 RFU in WT cells, slightly lower than uvrD by 250 RFU, Fig. 3 and Fig. S2). This indicated that the SOS response was active in some cells. By 6 hours, the majority of the cell population were seen to display a red fluorescence signal, and correspondingly an RFU of 7300 was detected using the plate reader (similar to *uvrD)*. The fluorescence detected at 24 hours post fork-blockage had increased dramatically (22,000 RFU) though plasmid loss and cell death were observed alongside cells with increased mCherry intensity (Fig. 3). This result was similar to the *uvrD* 24-hour sample (Fig. S2). As expected, over the time course, no mCherry fluorescence was seen in any *recA* cells by microscopy (data not shown), and the plate reader detected under 1800 RFU of mCherry (Fig. S2), confirming that the SOS response is not induced in the absence of RecA. The effect upon SOS induction of releasing the replication roadblock by the addition of AT after 2 hours of fork blockage was also assessed. No observable SOS induction was seen 2, 4 or 22 hours post AT addition in either WT, uvrD- or recA-strains (Fig. 3 and Fig. S2). This confirms that the SOS response is not induced by production of the replication roadblock within the first 2 hours, and thus the initial replication fork processing events observed are not influenced by the SOS response.

Replication fork processing and RFR can be induced across the population by a shift to the non-permissive temperature for the dnaBts strain, and this is observed to occur rapidly (44). Therefore, DnaBts can be utilised to collapse forks and monitor RFR events over time in a population of cells. We cannot rule out the possibility that prolonged exposure of *dnaBts* cells to 42°C following the establishment of the replication fork block will induce SOS. However, processing events seen at the early time points after 90 minutes of inducing the replication roadblock will not be influenced by SOS induction.

### RFR is more dependent on RecQ than RecG or RuvABC

To determine the contributions of various proteins to RFR at a nucleoprotein block *in vivo*, mutations in *recG, ruvABC* or *recQ* were introduced in the *dnaBts* strain containing the *tetO* array. Once the formation of a replication blockage had been verified by fluorescence microscopy, the cells were shifted from 30°C to 42°C to inactivate DnaBts. The previous experiment established that a strain carrying DnaBts had the majority of its blocked forks reversed out of the array within an hour at the non-permissive temperature (Fig. 2). To more closely investigate the timing of RFR, samples were taken at 15-minute intervals in both the dnaBts and dnaBts + helicase mutant strains at 42°C and the DNA structures present in the array region were analysed via 2-D gels. If the Y-structured DNA indicative of a blocked replication fork persists at the array region within one of the strains, it indicates that the mutation is affecting the cell’s ability to process the replication fork. The region upstream of the array was also visualised to see if the observed levels of Y-structures at the array correlated with the presence or absence of upstream DNA structures. In dnaBts cells with replication blocked by addition of arabinose, a shift to 42°C resulted in a drastic loss in Y-structured DNA signal at the 15 minute time point; 15% Y-shaped DNA remained from the 50% seen initially (Fig. 4B). The DNA upstream was mostly linear with a low level of Y and HJ DNA, and remained so over the hour (Fig. S1, Fig. 2C).

**Fig 4:**
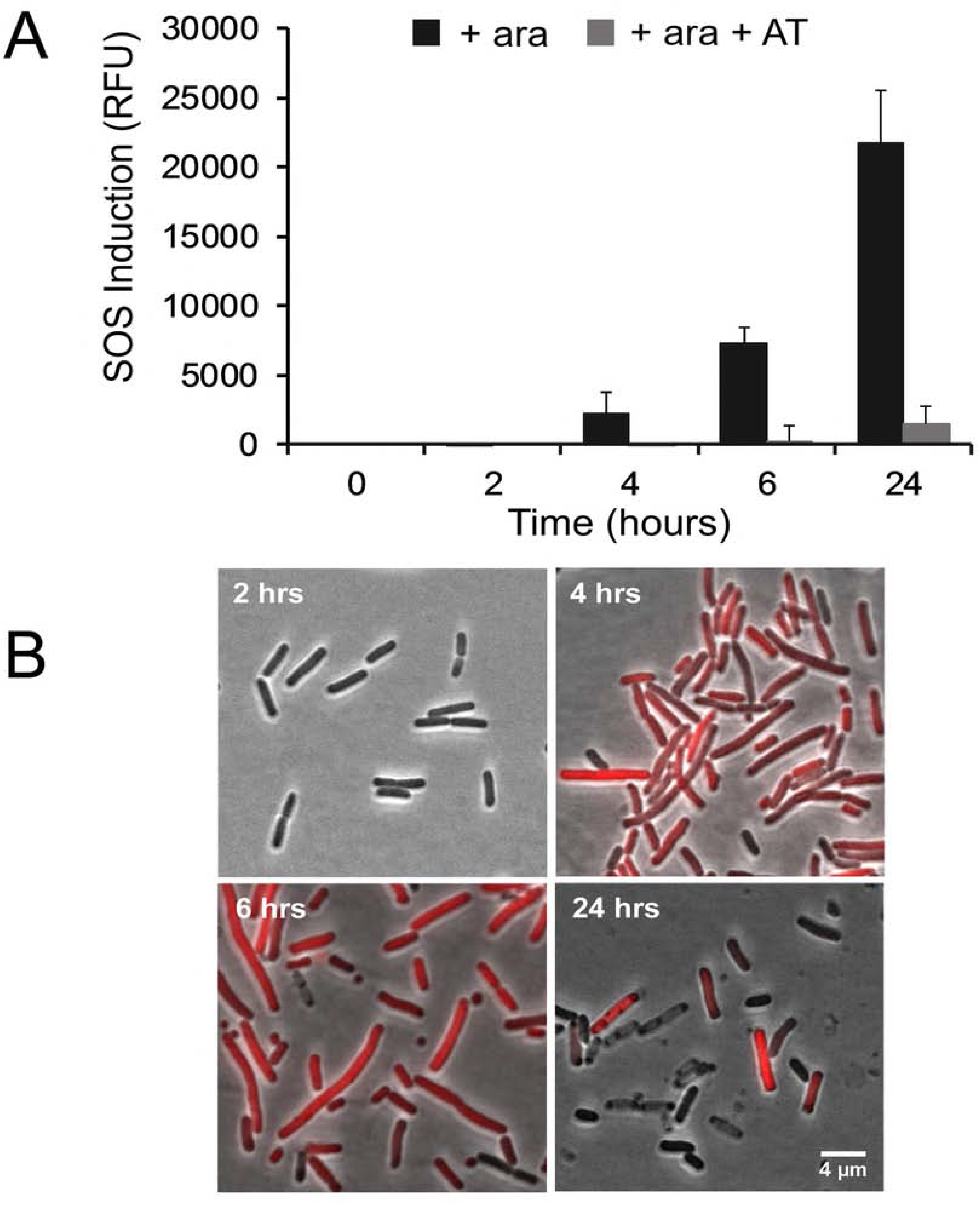
RFR frequency is reduced in *dnaBts recG, dnaBts ruvABC* and *dnaBts recQ* mutants. 2-D gel analysis of replication block (+ara) after 0, 15 and 60 minutes at 42°C (the non-permissive temperature for DnaBts), following an initial growth with arabinose at 30°C for 2 hours. (*A*) DNA structures within the array region. (*B*) Percentage of Y-structured DNA within the 5.8 kb array region.^#^ Denotes significant difference between Y-DNA amount in dnaBts and the mutant at 0 min (*P < 0.05*). At 15 and 60 min, the percentage of Y-DNA remaining in each mutant was determined and deemed significantly different to that remaining in dnaBts (* P <0.05, ** P < 0.01, *** *P* < 0.001) (*C*) Replication intermediates detected in the region upstream of the array. n = 3

Following the same procedure to block DNA replication in dnaBts recG cells, a strong Y-shaped forked DNA signal was observed indicative of the replication roadblock, but at a significantly lower amount than *dnaBts* (Fig. 4A and B). Upstream of the array a strong HJ signal is visible, and is reproducibly stronger than in other genetic backgrounds (Fig. 4C). At the permissive temperature for DnaBts, the steady turnover of forks and their processing and subsequent restart should be occurring, yet without RecG there is a marked accumulation of unresolved HJ signal, indicating a role for RecG in the processing of these HJs. Shifting the dnaBts recG cells to 42°C resulted in a reduction in Y-DNA signal within the array region as fork-collapse occurs to an extent similar to that seen in WT (Fig. 4A); after 15 minutes, 20% of the signal was present as forked DNA in the array, which further reduced to 13% by an hour (Fig. 4B), levels that are slightly higher than seen for *dnaBts*. This was accompanied by the loss of HJ and Y-shaped signals upstream over the 42°C time-course (Fig. 4C), but this occurred more slowly than in *dnaBts* where by 60 minutes almost all the signal had disappeared (Fig. S1). It is plausible that the slightly reduced level of Y-shaped DNA seen at the array upon replication blockage is a result of a subtle alteration in the equilibrium between blocked replication forks at the array, and the collapsed forks undergoing processing. The higher level of HJ seen may reflect that RFR takes longer to be resolved in a *recG* mutant with DNA existing as a HJ for a longer time, even in the presence of RuvABC. It is noteworthy that the position of the HJ signal in the 2-D gels is that of a so-called “X spike”, which is distinctive of a HJ made from two dsDNAs of full length. If degradation on one arm had occurred, or if the HJ was formed by homologous recombination using a broken strand within this region, then it would migrate in the “cone signal” area and not as seen. The HJ is, therefore, distinctive of a RFR event.

The *dnaBts ruvABC* mutant produced a replication block at a similar level to *dnaBts* at 30°C (Fig. 4A). Inducing fork collapse at 42°C led to a rapid drop in Y-DNA signal as seen in *dnaBts* (Fig. 4B). However, after an hour at 42°C a slightly higher percentage of DNA remained as a fork structure, 20% compared to 12% for *dnaB*ts. The prominent HJ seen upstream in the *recG* mutant was not present in *dnaBts ruvABC* and the non-linear DNA structures that were present were largely processed in the course of the hour at 42°C, similar to dnaBts (Fig. 4C; S1). It is noteworthy that this processing of the upstream DNA appeared efficient even in the absence of RuvABC.

The greatest persistence of Y-shaped DNA following a shift to 42°C was seen in the dnaBts recQ mutant. Upon establishment of a replication block at 30°C, 65% of DNA was Y-shaped, a significantly larger proportion than in *dnaBts* (Fig. 4A and B). After 15 minutes at 42°C the DNA remained largely in the forked-DNA structure (Fig. 4A). Quantitatively, forked DNA now made up 44% of the total DNA, signifying that only a third of the initial Y-shaped DNA had been processed. The fork signal persisted even after an hour at 42°C, with 33% of the total DNA still being in a Y-structure (Fig. 4A and B) i.e. roughly half the initial Y-shaped DNA remained after 1 hour, compared to 23% of initial forks in *dnaBts* (Fig. 2B). Upstream of the replication block processing intermediates were present at all times; this could reflect that processing of the upstream DNA was also slower in recQ mutants, or that there was a constant low level of processing of the Y-shaped structures to generate more upstream signals that continued over the full hour (Fig. 4C). The slowed processing of the stalled fork could explain why a higher proportion of Y-structured DNA was initially seen at the array: it is the result of an alteration of the equilibrium between fork collapse/processing and restart similar to what was proposed for the RecG results above (Fig. 2B). The majority of processing of the Y-shaped DNA in these experiments occurs by RFR rather than direct nuclease cleavage of the Y-shaped DNA, as RecQ has no nuclease function. Following inactivation of DnaBts it has been seen that both ExoI and RecJ can contribute to degradation of the nascent DNA strands (50), and RecJ activity has been shown to be stimulated by RecQ activity that provides a suitable substrate (51). Therefore, inactivation of RecQ may inhibit processing that leads both to RFR and exonuclease digestion to yield Y-shaped structures upstream.

These findings demonstrated that *recG, ruvABC* and *recQ* mutants exhibit a diminished ability to reverse replication forks, implying that all three proteins are involved in RFR. It also implies that very little direct endonuclease digestion of the Y-shaped DNA occurs, as it is difficult to envisage this being restored to a stable Y-structure at the non-permissive temperature for DnaBts. However, the most extensive deficiency to RFR was seen in *dnaBts recQ* indicating that RecQ is key to the majority of RFR under these conditions.

### RecG and RuvABC act in a distinct pathway from RecQ to process stalled forks

To determine whether RecG, RuvABC and RecQ act synergistically to perform RFR, we investigated strains with combinations of these gene knockouts. If these proteins act independently, the absence of multiple proteins should be an additive effect, and the signal indicative of forked DNA that accumulated in the array region during induction of the replication blockage will persist at a higher proportion than in any of the single mutants alone upon DnaBts inactivation.

In the *dnaBts recGrecQ* mutant, the proportion of forked-DNA at the array was significantly higher than in a *dnaB*ts mutant; it remained at a high intensity in the array region 15 minutes after the shift to 42°C when it reduced to 37% from 64% (Fig. 5A and B). The intensity of this Y-DNA signal was unchanged at the 60-minute time point (37%, Fig. 5A and B). Therefore, 57% of the initial forks within the array region failed to be processed in the dnaBts recGrecQ mutant even after one hour, marginally more impaired than a recQ mutant. The HJs and Y-arcs detected upstream after the initial replication block (0 minutes) in *dnaBts recGrecQ* mutant included a prominent HJ signal, like those seen in a recQ mutant. Shifting the cells to 42°C resulted in a loss of the Y-DNA arc and a decrease in the HJ signal after 60 minutes, although they were still very prominent at 15 minutes (Fig. 5C).

**Fig 5:**
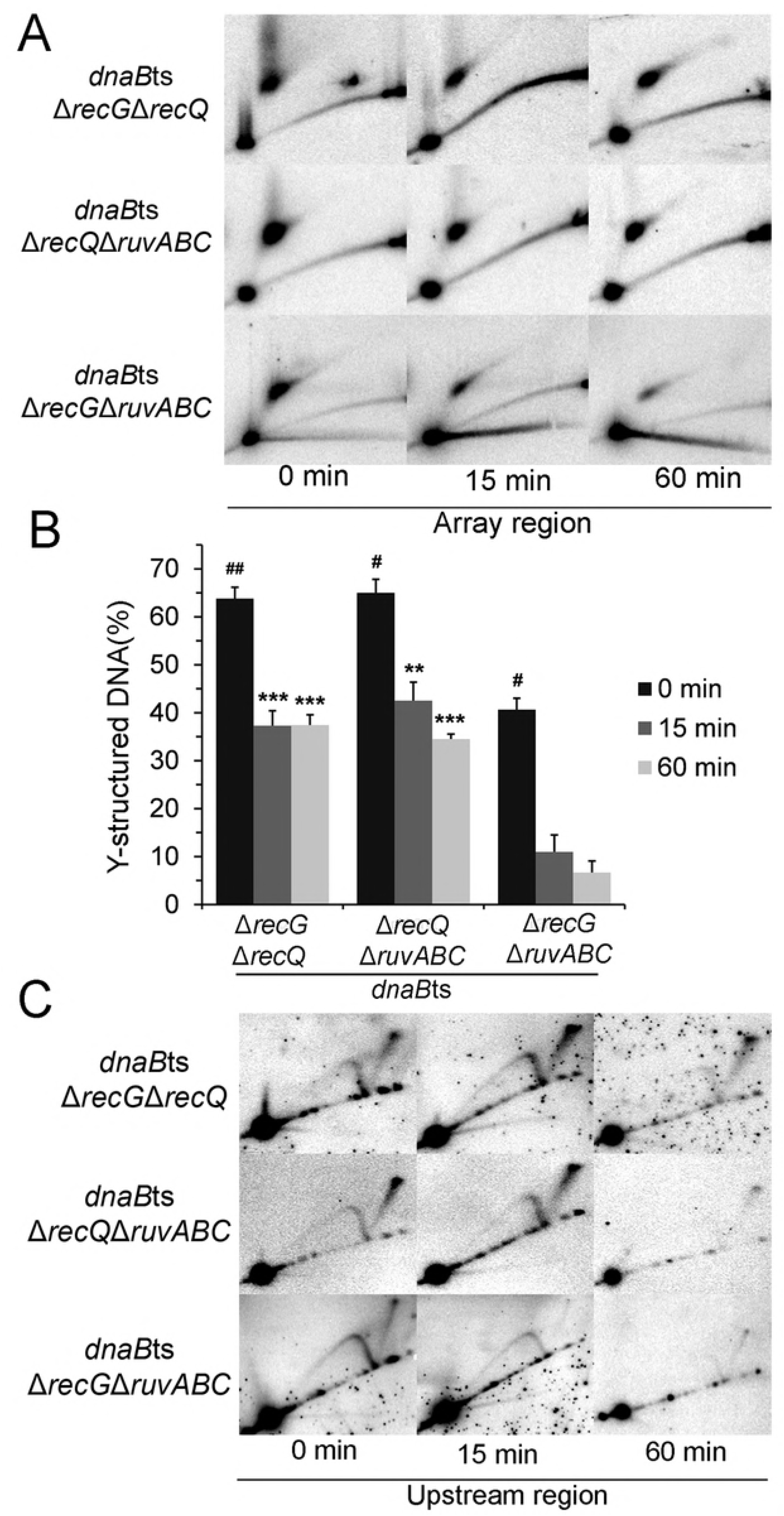
RFR in *dnaBts recGrecQ, dnaBts recQruvABC* and *dnaBts recGruvABC* strains over time. 2-D gel analysis of replication block (+ ara) after 0, 15 and 60 minutes at 42°C (the non-permissive temperature for DnaBts), following an initial growth with arabinose at 30°C for 2 hours. (*A*) DNA structures within the array region. (*B*) Percentage of Y-structured DNA within the 5.8 kb array region.^#^ Denotes significant difference between Y-DNA amount in dnaBts and the mutant at 0 min (^#^ *P* < 0.05,^##^ *P* < 0.01). At 15 and 60 min, the percentage of Y-DNA remaining in each mutant was determined and deemed significantly different to that remaining in dnaBts (** P < 0.01, *** *P* < 0.001) (C) Replication intermediates detected in the region upstream of the array. n = 2–3

Analysis of the dnaBts recQruvABC strain bore a close resemblance to that of dnaBts recGrecQ and dnaBts recQ. Once shifted to 42°C the intensity of the forked DNA signal remained relatively high at both the 15- and 60-minute marks (Fig. 5A). 43% of the total DNA remained forked at the block site after 15 minutes with a small decrease to 35% by 60 minutes. Therefore, of the original Y-shaped signal 54% remained at 60 minutes (Fig. 5B). Upstream DNA signals were also akin to those seen in the dnaBts recQrecG mutant, as they were maintained at a high intensity upstream of the block once fork collapse was initiated and observed at 15 minutes, but became fainter after an hour (Fig. 5C). The similarity in the signals seen in both *recGrecQ* and *recQruvABC* could mean that that RecG and RuvABC contribute equally to RFR, or could act in the same pathway together. However, RecQ’s contribution to RFR appears to far outweigh either RecG or RuvABC.

When dnaBts recGruvABC cells were grown at 30°C and the replication roadblock was induced, ∼45% of the total DNA was seen to be Y-shaped. This percentage is similar to that seen with dnaBts recG (46%), and is lower than seen with other strains, possibly reflecting a longer lived HJ intermediate in the absence of RecG. Upon shift to 42°C the signal strength of the forked DNA decreased to ∼10% at 15 minutes and by an hour a further decline to 7% Y-DNA occurred (Fig. 5A and B). The non-linear DNA structures detected in the dnaBts recGruvABC mutant upstream of the block dissipated over time becoming faint after 15 minutes and almost disappeared by 60 minutes (Fig. 5C). The signal intensity seen in this region more closely resembled that of dnaBts recG and dnaBts *ruvABC* rather than maintaining the stronger signals of dnaBts recGrecQ and dnaBts recQruvABC. Both RecG and RuvABC have several reported functions, some of which overlap. The 2-D gel profiles of the double recGruvABC mutant closely resembled that of each single mutant, suggesting that they did not have an additive effect, and may be working in the same pathway.

Finally, the effect of the absence of all three proteins were analysed collectively via a *dnaBts recGrecQruvABC* mutant. Based on the previous trends, it was expected that the triple deletion mutant should resemble either double mutant containing *recQ*. Indeed, 60% of the initial Y-shaped DNA signal remained after 60 minutes at 42°C (Fig. 6). Additionally, the signal patterns detected upstream over the hour of fork collapse in the triple mutant were consistent with the single and double mutants, where the non-linear DNA signals were strong at 0 and 15 minutes but reduced by an hour (Fig. 6). This again verified that RecQ contributes greatly to RFR.

**Fig 6:**
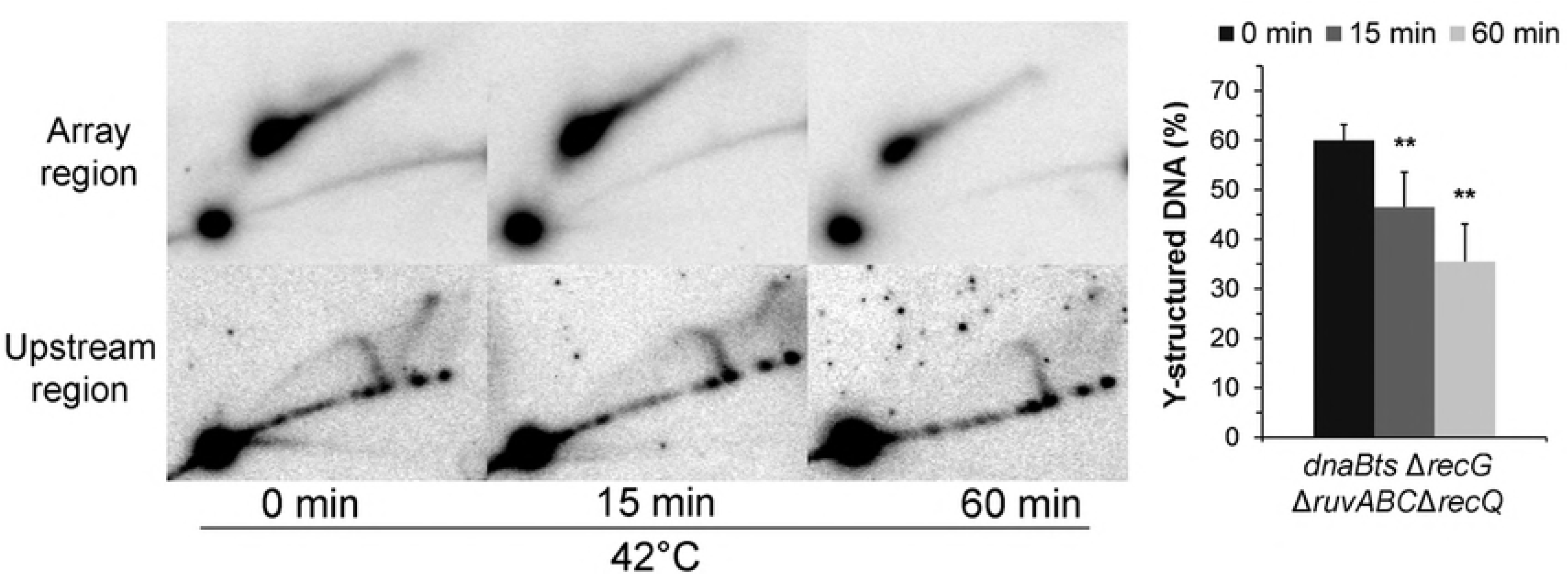
RFR in a *dnaBts recGruvABCrecQ* mutant. 2-D gels of the array and upstream regions following 0, 15 and 60 minutes of replication blockage at 42°C (after an initial growth with ara at 30°C for 2 hours) alongside a graph of the percentage of DNA in the forked structure at each time point. At 15 and 60 min, the percentage of Y-DNA remaining in each mutant was determined and deemed significantly different to that remaining in dnaBts (** P < 0.01). n = 2

### Restart of replication is impaired in a *recG* mutant

It has been previously established that the TetR-YFP replication roadblock can be relieved with the addition of AT, resulting in the restart of replication (Fig. 1). The addition of AT for 10 minutes following 2 hours of replication roadblock restored viability to levels seen in isogenic cells that never had arabinose to induce replication stalling. 2-D gel analysis of this replication resumption in *dnaBts* saw the disappearance of forked DNA (Fig. 7), leaving solely linear DNA, in both the upstream and array regions after 10 minutes AT exposure, which correlates with the restoration of viability (Fig. 1). The absence of forked DNA meant that all the intermediates previously seen (Fig. 2) had been processed within this 10 minute period. The most likely explanation is that replication restart had occurred across the population of cells. Importantly, once the cells had undergone the temperature shift to 42°C for one hour and then back to 30°C to allow re-establishment of the replisome, again only linear DNA resided in the 2-D gels after 10 minutes AT exposure (Fig. 7A) (compare to signals in Fig. 2 where AT was not added). As previously noted (Fig. 1) the counting of cell numbers with multiple foci for *dnaBts* revealed that there was a slower release of the block under these conditions, however, this delay wasn’t seen in the 2-D gels. This suggests that the replication forks have restarted within 10 minutes and the array has been copied, but there has not been sufficient time for the two daughter copies to segregate from each other.

**Fig 7:**
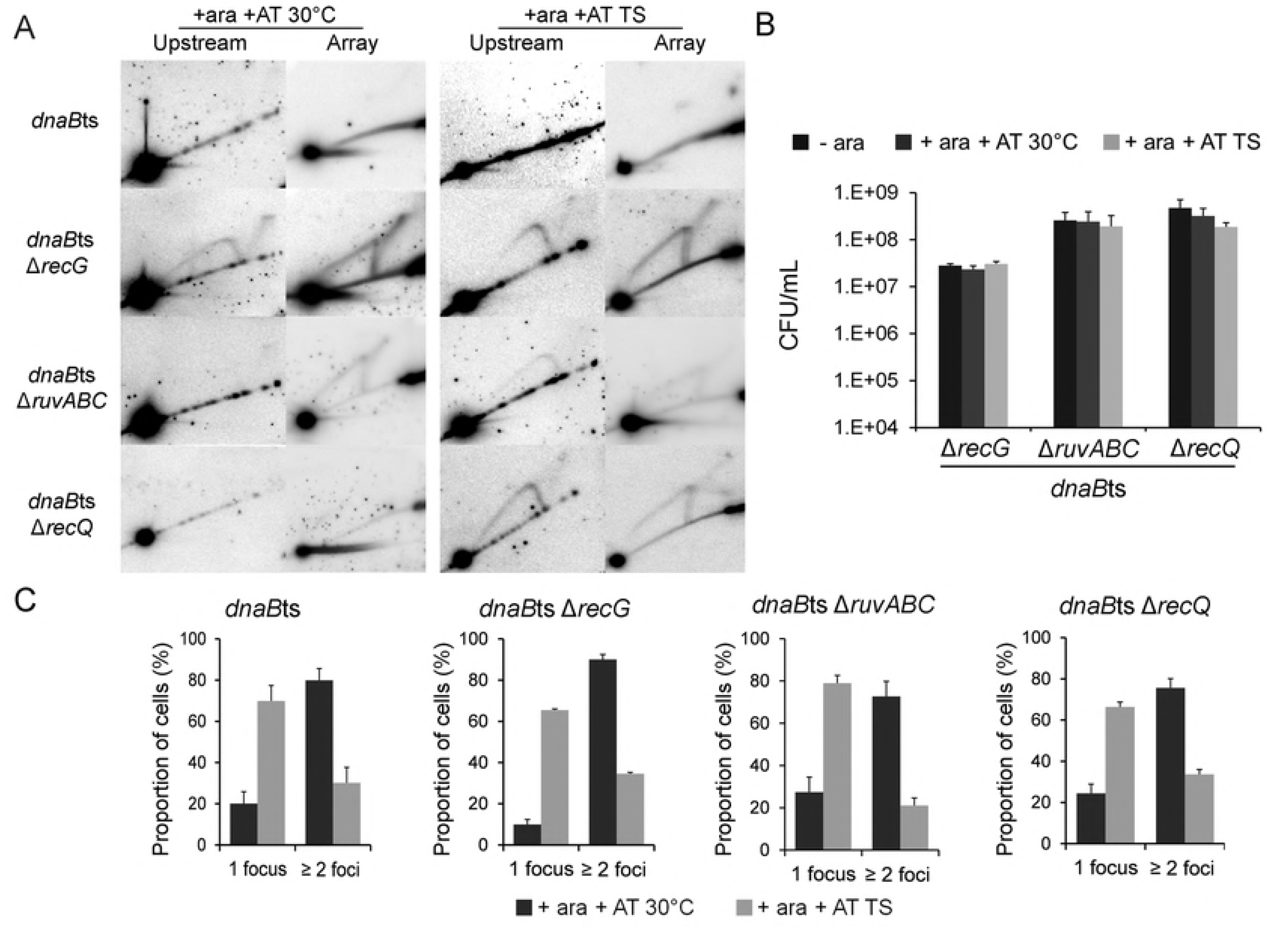
Replication restart is severely impaired in a *dnaBts recG* mutant and slightly impaired in *dnaBts ruvABC* and *dnaBts recQ* mutants. Subsequent to an initial growth at 30°C with ara, a subset of cells were treated with AT for 10 minutes to release the block (+ara +AT 30°C). The remaining cells underwent a temperature shift from the 30°C to 42°C then returned to 30°C (TS) and exposed to AT for 10 minutes (+ ara + AT TS). Replication intermediate structures visualised by 2-D gel analysis in the array and upstream regions 10 minute after addition of AT (*A*), cell viability (*B*) and foci percentages (*C*) under indicated conditions are shown. See also Fig. S3 and Fig. S4.

The *dnaBts recG* strain produced extremely prominent HJ and Y-shaped signals upstream of the replication roadblock (Fig. 4C). The ability of these cells to restart replication was examined by removal of the TetR block by addition of AT for 10 minutes. It was seen that substantial levels of Y-shaped DNA and HJ were present both at 30°C and after temperature shift to 42°C (1 hour) and back to 30°C, in both the array and upstream regions (Fig. 7A). However, the former spot on the Y-arc, indicative of replication blockage, is now absent showing successful release of the protein roadblock. The absence of RecG left a proportion of HJs and forks that were yet to be processed and/or migrated in order to restart replication in a timely manner. Despite the inability of the cells to resolve these intermediates within the 10 minute window, they were able to eventually recover their viability upon release of the block (Fig. 7B), indicating that this was a delay rather than a failure to restart DNA replication. To ensure the DNA structures visible in the 2-D gels were not an artefact of the presence of DnaBts, the same assay was performed on a recG strain with WT DnaB (Fig. S3). The resulting 2-D gel showed similar signals as for the dnaBts equivalent.

The addition of AT to the dnaBts ruvABC mutant resulted in barely visible HJ and Y-arc signals upstream of the array, both at 30°C and after temperature shifting. In the array region, the HJ and Y-arc signal were faint, but clearly present, though at a lower level than in the equivalent *recG* mutant (Fig. 7A). This may also suggest a slight delay in replication restart, and longer persistence of HJ intermediates in the absence of RuvABC. Overall, the majority of HJs were able to be processed or resolved in the absence of RuvABC (compare Fig. 4 to Fig. 7). Following addition of AT to allow replication to proceed through the array, the number of foci within each cell was similar to that seen in dnaBts; 72% of cells had two or more foci at 30°C but following the temperature shift to 42°C only 21% showed multiple foci in the presence of AT (Fig. 7C). Cell viability was also completely restored by 10 minutes treatment with AT (Fig. 7B).

Linear DNA was the only DNA structure visualised by 2-D gels in both the array and upstream regions of a dnaBts *recQ* mutant following the release of the roadblock via AT addition at 30°C (Fig. 7A). Replication restart was also seen to be similar to *dnaBts* in terms of cell viability and the number of foci observed per cell as replication restarted (Fig. 7B and C). However, following the temperature shift, lingering intermediate signals remained in both regions (Fig. 7A). Cell viability was seen to be fully restored in the dnaBts recQ following addition of AT (Fig. 7B) suggesting replication restart does eventually occur across the population. Furthermore, the double and triple mutants did not affect replication restart, as their viabilities were shown to be restored to levels not significantly different from untreated (Fig. S4).

## Discussion

It has been proposed that almost all replication forks in *E. coli* will encounter DNA damage on the template during replication in normal growth conditions (5), and that collision with nucleoprotein complexes provides an even greater impediment to replication progression than DNA damage (3). This study examined the relative contributions of RecG, RuvABC and RecQ in the reversal and processing of replication forks that had collapsed at a nucleoprotein block, in vivo. An array of repressor-operator complexes allowed replication to be arrested at a specific location on the chromosome. Introduction of a temperature sensitive allele of DnaB enabled the synchronous deactivation (and possible dissociation) of the replisome across the cell population in order to be able to assess the DNA processing events that followed. Processing of the stalled replication fork DNA can be seen by the loss of Y-shaped DNA at the block upon replication fork collapse, concurrent with HJ being seen in the region upstream of the array. The simplest conclusion is that collapse of the replisome leads to RFR, the first processing event thought to occur at a stalled fork. RecG and RuvABC, two branched-DNA specific motor proteins, have previously been implicated in the reversal of stalled replication forks (8, 20). In this study, both *dnaBts recG* and *dnaBts ruvABC* mutants showed a similar deficiency in fork processing upon replisome dissociation, with ∼30% and ∼35% respectively of the initial forked DNA at the block site remaining after an hour at 42°C, compared to ∼24% in *dnaBts*. This increase in the quantity of forks remaining at the block lead us to conclude that both RecG and RuvABC assist in RFR of a blocked fork. However, a greater percentage of forks were found to remain at the block in the recQ mutant (50%) highlighting the key role of RecQ in facilitating RFR.

Previously, RecQ has been shown to bind and unwind a diverse range of DNA substrates in vitro (36), including unwinding of dsDNA without a ssDNA gap (52), and displays a 3’–5’ movement along one strand. The RecQ catalysed unwinding is optimal when two or more monomers act on a single DNA substrate, providing an increased rate of unwinding. This in vitro mechanism of action for RecQ is consistent with the in vivo evidence presented here of RecQ’s involvement in RFR. The involvement of RecQ in the RecFOR-mediated homologous recombination pathway suggests that it migrates on one strand to produce or enlarge ssDNA gaps. RecJ may then also aid the process by degrading the displaced strand from its 5’ end and enabling RecFOR-mediated loading of RecA onto the ssDNA to initiate strand exchange. At a stalled fork, RecQ could load onto the lagging strand gap and act to enlarge the ssDNA region. RecA may then mediate strand exchange to re-pair the two template DNA strands, leading to fork reversal (Fig. 8). Alternatively, RecQ may play another, not mutually exclusive, role by acting at a fork which has been reversed by RecA-mediated strand exchange. Two RecQ proteins proceeding in opposite directions from the branch point of the fork along the lagging strand template (unpairing the nascent lagging strand) and the nascent leading strand (unpairing it from the leading strand template) could directly facilitate RFR by converting both nascent strands into ssDNA allowing their mutual pairing.

**Fig 8:**
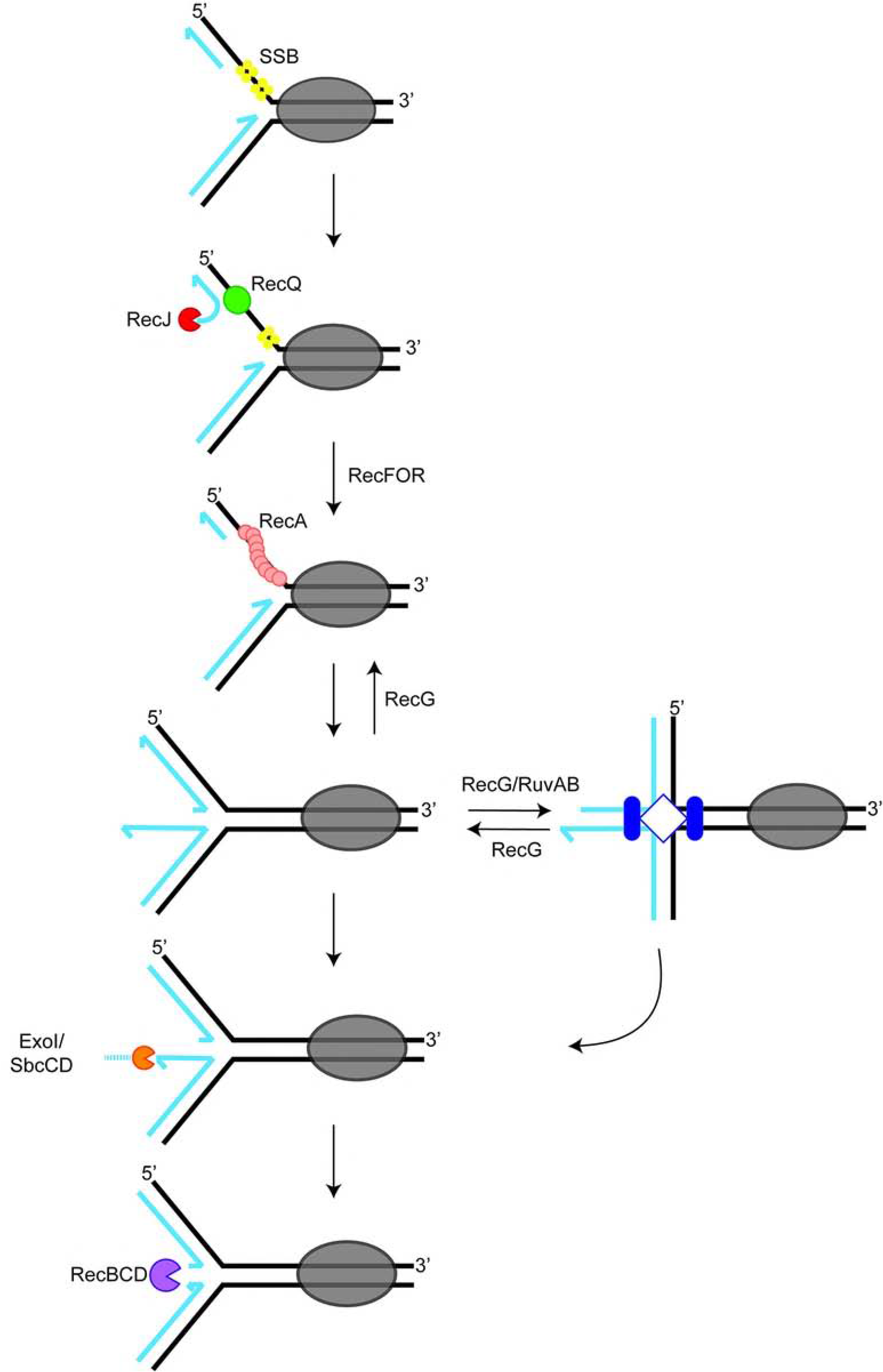
Model for replication fork reversal at a protein roadblock. Template DNA is shown as black lines and newly replicated strands are in blue. A proteinaceous roadblock to replication is shown as a grey oval. The most likely disposition of the leading and lagging strands is shown, with the leading strand coming close to the site of the block. Upon this substrate RecQ can act, possibly in concert with RecJ, to lengthen the ssDNA gap on the lagging strand template. RecFOR could then load RecA on the ssDNA, leading to strand exchange which pairs the two template strands, displacing the nascent leading strand. The initially regressed fork can then be acted on by a number of pathways such as the known branch migration proteins RecG and RuvAB which could move the HJ upstream. RecG may also reverse this process to re-create the Y-shaped DNA to allow replisome reloading. Nuclease action by ExoI/SbcCD, RecQ/J and/or RecBCD could also remove the two nascent DNA strands to eventually re-generate a Y-shaped fork upstream from the site of the roadblock.

The key role for RecQ found in this study reflects evidence on replication fork processing in eukaryotes. The human RecQ homologs BLM, WRN and RECQ5 have been heavily implicated in regressing replication forks (40, 41, 53–55), and their absence leads to a predisposition to cancer, genetic instability and premature aging. In yeast the Sgs1 protein is the single RecQ homologue and has been show to associate with stalled replication forks following HU-treatment (56). Thus, the proposed role of RecQ in RFR in *E. coli* may be functionally conserved between bacteria and eukaryotes.

RecG has been shown *in vitro* to be able to regress Y-shaped forked DNA into a HJ, but it has a preference for forks where the leading strand is absent or there is a gap on the leading strand (57). However, it would seem likely that a replisome blocked by a protein on the DNA template would synthesise DNA on the leading strand at least as far as on the lagging strand, or further. Blockage of replication at the Tus/*ter* complex resulted in leading strand synthesis right up to 4 bases from the conserved GC6 of the *ter* site both *in vivo* and *in vitro*, whereas lagging strand synthesis only approached to 50–100 nucleotides from the bound Tus protein (58, 59). A similar disposition of strands would be expected for any protein block on the DNA template. Furthermore, the crystal structure of RecG on a model forked DNA structure showed that it binds to the duplex DNA of the template ahead of the stalled fork in order to catalyse regression (26). Although the reported structure only contained 10 bp of template DNA ahead of the branch point, domain 3 of RecG is positioned to contact DNA over a further region of approximately 10–15 bp. If another protein were present blocking the replication fork then it seems likely that this would sterically hinder access of RecG to the 20–25 bp of DNA it requires for binding, preventing it from catalysing fork reversal. Together, the structural preference of RecG and the steric hindrance from a protein roadblock may explain why RecG appeared to play a relatively minor role in RFR following encounter of the replisome with a protein roadblock. RecG may be more active in RFR on other substrates.

RuvAB has been shown to be able to regress a fork *in vitro* to form a HJ, but it does so with low efficiency, preferring to unwind DNA in the opposite direction from that required to form a HJ (60). RuvAB has instead been shown to efficiently migrate a pre-formed HJ *in vitro* (61). Because of this, RecG has been suggested to first regress the stalled fork to form a HJ which RuvAB subsequently branch migrates (29). Additionally, RuvC has been shown to cleave the HJs formed by RecG regression (19). Consistent with this, a relatively minor role for RuvABC in direct RFR was observed in this study. The similarity between the phenotypes of *ruvABC, recG* and the combination recGruvABC double mutant in these assays, supports the notion that they may play roles in the same pathway. However, the majority of RFR observed here was dependent upon the action of RecQ. It is likely that the pathway utilised for processing a stalled fork is dependent upon the exact structure of the DNA at the branch point, namely the relative disposition of the leading and lagging strands. A heterogeneous population of DNA structures in vivo may account for the fact that a proportion of forks are processed in the absence of RecQ. In addition, multiple overlapping pathways may exist to catalyse RFR and in the absence of RecQ a less efficient pathway predominates. The diversity in the possible structures of the DNA at a replication block may account for differences seen in processing pathways utilised by the cell, particularly after treatment with UV where previously a RecF-pathway dependent processing event has been found to repair the fork (62).

When mutations in recQ were combined with recG and ruvABC, it was found that there was a further decrease in the efficiency of RFR with ∼60% and ∼55% of replication forks, respectively, remaining unprocessed after one hour following inactivation of DnaBts. The recQrecGruvABC mutant showed a similar decreased efficiency of RFR to the double mutants, with ∼60% of the replication forks remaining after 1 hour at 42°C. This result again suggests that RecG and RuvABC may be associated in the same processing pathway. Concurrently with this there was an increased HJ signal upstream of the roadblock in the double mutants, suggesting a delay in dealing with processing the HJs that result from the reduced level of RFR.

RFR is only one possible mechanism to deal with a collapsed replication fork; another proposal is that direct endonuclease action on an arrested fork can lead to breakage of one arm at the fork, which can then be repaired by homologous recombination. However, the results presented here show that this did not occur at the majority of forks. Stalled Y-shaped DNA in a recQ mutant was largely stable over an hour suggesting nuclease action must be slow or only works on a sub-population of fork structures, with the majority normally being processed by RFR. Upon cleavage, the released DNA arm would most likely be acted upon by exonucleases such as RecBCD prior to loading of RecA and the initiation of strand exchange. Homologous recombination initiated by RecA would produce a HJ that RuvC would be required to resolve in order to restore the Y-shaped fork structure. Efficient restart and recovery of viability in a *ruvABC* mutant argues against this pathway being utilised to any great extent. Further, in the absence of functional DnaB (and hence the absence of the entire replisome on DNA) the Y-shaped structure resulting from homologous recombination would not be located at the site of the original blockage, but the branch of the Y would be located further upstream depending on the extent of exonuclease processing prior to loading of RecA. This is inconsistent with the long-lived Y-signal in the *recQ* mutants at the site of the roadblock. Therefore, we conclude that once replication is blocked by a protein complex on DNA it is mostly processed by RFR, and that this occurs rapidly (44). For the same reasons it is most likely that the HJ observed in the DNA immediately upstream of the replication roadblock is the direct result of RFR, rather than being an intermediate in repair of a free double stranded end by homologous recombination. Indeed, mechanistically it makes sense for the cell to avoid producing a broken DNA molecule if possible, as this can lead to potentially mutagenic repair processes.

The other possible processing event at the collapsed replication fork is the action of exonucleases that digest the two nascent DNA strands, an activity that has been previously reported when DnaBts is inactivated and is attributed to RecJ and ExoI (50). There is indeed a Y-arc signal seen in the upstream DNA along with HJ that could be the result of exonuclease action. However, what is unclear is whether the exonucleases act on the DNA of the Y-shaped fork, or on the ends of the reversed HJ produced by RFR, as these would both produce the same structure in the upstream DNA. It should also be noted that the HJs detected are most likely an underestimate of the HJ levels *in vivo*. Each HJ is free to branch migrate and should be fully homologous throughout its sequence. If branch migration occurs to move the branch point all the way to one end of a DNA arm, then the HJ can dissolve into two DNA duplexes that would migrate in the linear spot seen. This process would be essentially irreversible. Cell lysis, washing and DNA digestion occur over the period of days giving ample time for some HJ dissolution. Similarly, if one arm of the HJ has been partially digested by exonucleases, then branch migration to the end of this arm would convert the HJ to Y-shaped DNA. Given these caveats about the observed level of HJ seen, then it seems plausible that RFR is the major pathway utilized in cells and the HJ lvel observed underestimates this.

The SOS regulatory response is induced by the prolonged presence of RecA filaments on ssDNA, and may be induced during long periods of fork stalling and processing. An exact time scale for SOS induction from fork stalls cannot be universal as it will likely vary between cells and the pathway for DNA processing that each chooses. To detect SOS induction, a reporter strain was created to determine the activation time of SOS response once replication forks had been stalled in our system. Importantly, no evidence of SOS induction was detected 2-hours after creation of a replication roadblock (Figure 3). During the experiments described here the typical fork-blocking induction period is 90 minutes, followed by an additional hour, at most (at 42°C or 30°C), leading one to conclude that our artificial block does not induce the SOS response in WT cells. The *dnaBts* strain behaved similar to WT, with the *dnaBts* allele used simply as a means to amplify processing events and the majority of RFR occurring in the first 15 minutes (Figure 2). Thus, the RFR visualised in a *dna*B*ts* strain remains a valid interpretation of the mutants’ results and not an artifact of SOS response processing within the timeframe assessed. The SOS response begins to activate minimally only after 4 hours of fork stalling, exacerbated to the entirety of the cell population after 24 hours. The release of a replication fork block by addition of AT did not lead to activation of the SOS response suggesting that replication can recover quickly once TetR-YFP leaves the DNA. This result also rules out the possibility that the mCherry protein did not have sufficient time for folding within the 2 hour replication block- no fluorescence was seen in these cells at longer time points either.

Although the effect of deletion of recG upon RFR was modest, it was clear that in the absence of RecG a much stronger HJ signal was seen in the region upstream of the replication roadblock. This implicates RecG in the timely processing of HJs. The simplest and least recombinogenic mechanism to deal with a HJ produced by RFR is to branch migrate the junction back into a Y-shaped fork, upon which the replisome can be reloaded, and this may be a role played by RecG. *recG* mutants also showed an accumulation of HJ structures during replication restart (Fig. 7 and Fig. S3) suggesting that HJs persist even after the replication roadblock has been removed in these cells, although the viability data suggests that the delay to recovery is not fatal and is eventually overcome. The deletion of ruvABC was also seen to increase the level and persistence of HJs in the upstream DNA region, consistent with the known role of the complex in branch migration and resolution of HJs.

It is noteworthy that HJs are seen to be present upstream of the replication roadblock in both recGruvABC and *recQrecGruvABC* backgrounds, and that the levels of these intermediates decline over time, along with a reduction in Y-shaped DNA at the block. There is clearly another process able to either migrate the HJs out of the region being probed or to resolve the HJ. As a result of this finding, we concluded that neither RecG nor RuvABC are absolutely required for HJs to be processed. RusA, the only other known endogenous HJ resolving enzyme in *E. coli*, is absent from the strains used in this study. Spontaneous branch migration of the HJ out of the region being examined could explain these results, but subsequent recovery of replication would then also depend upon the branch migration occurring to regenerate the Y-shaped structure to allow replisome reloading. An alternative pathway that has been proposed is the action of RecBCD (Fig. 8). One arm of the HJ formed by RFR has a DNA end. Single strand specific exonucleases can digest an overhang on this DNA end to produce a blunt ended duplex; ExoI and SbcCD can digest the 3’ overhang that would result from the leading strand having gone further than the lagging strand dsDNA end (63, 64). Conversely, RecJ can remove a 5’ overhang if the lagging strand was ahead of the leading strand upon RFR. Once the DNA end is blunt, RecBCD can load and processively digest both strands, gradually removing one entire arm of the HJ. The effect of this would be to convert HJ into a Y-shaped structure upon which replication proteins could be loaded allowing restart. It is notable that Y-shaped DNA is seen upstream of the replication roadblock when HJ is present, supporting this model of RecBCD action. RecBCD action can also lead to RecA loading after an encounter with a Chi site, eventually producing a second HJ by strand invasion. This double HJ can then be resolved either by RuvC or by branch migration and possibly action of a topoisomerase (such as Topo III) (65). It is also worth noting that RecQ could also be involved in digesting the free arm of the HJ produced by RFR; RecQ could act on a 3’ ssDNA overhang and subsequent translocation would eventually unwind duplex DNA allowing RecJ to digest the 5’ end strand (51). If the leading strand had proceeded further than the lagging strand then upon RFR the HJ would indeed have a 3’ ssDNA overhang.

The proposed model (Fig. 8) has some testable predictions. The establishment of the system described here to study replication fork processing in vivo can be built upon in future work to examine the roles of other proteins such as RecA, RecFOR, SbcCD, RecBCD, XonA and RecJ. This could lead to a comprehensive understanding of the complex and overlapping roles played by the many homologous recombination proteins in this key DNA repair process.

## Materials and Methods

### Strains and plasmids

Bacterial strains were derivatives of E. coli K12 AB1157 (66) which carry 240 copies of tetO in an array within the *lacZ* gene (67) and the pKM1 plasmid encoding the TetR-YFP repressor under the control of the P*ara* promotor (44). Lambda Red recombination was conducted initially to replace *recQ, recG*, *ruvABC, recA* and *uvrD* with a spectinomycin resistance cassette (68). The resistance cassette was subsequently removed by Flp recombinase, leaving only an FRT site flanked by the start and stop codons (68). The temperature-sensitive dnaBts allele (69) and all gene knockout strains were created by P1 transduction and confirmed by PCR (Table S1 for list of strains). The *sulAp*-mCherry construct, created by fusing the *sulA* promotor to mCherry, was inserted into the chromosome approximately halfway between ori and ter on the left replichore by Lambda Red recombination in a non-coding region while the constitutive *sulA* gene remained.

### Bacterial growth

Cultures grown overnight at 30°C in L-broth were diluted to OD_600nm_ = 0.01 in a dilute complex medium (0.1% tryptone, 0.05% yeast extract, 0.1% NaCl, 0.17 M KH_2_PO_4_, 0.72 M K_2_HPO_4_). Cells were supplemented with ampicillin (100 µg/ml) and/or kanamycin (50 µg/ml) as required. Production of the fluorescent repressor TetR-YFP was induced by addition of 0.1% (w/v) arabinose once cells had reached at least OD_600nm_ = 0.05. Cells were then incubated for 90 minutes and examined under a fluorescence microscope as described previously (70) to confirm the extent of replication blockage across the cell population. A minimum of 200 cells were assessed and foci counted. Cells were shifted to 42°C for an hour to induce replisome collapse in the temperature sensitive *dnaBts* strain. Subsequently, the cells were shifted back to 30°C for 10 minutes. Tight repressor binding was relieved by the addition of the gratuitous inducer anhydrotetracycline (AT; 10 µg/ml). Cell viability was determined by performing a 10-fold serial dilution and 5 µl of each dilution was spotted on agar containing the required antibiotic (and AT if needed). Selected dilutions were spread onto agar to determine CFU/ml. All plates were grown at 30°C overnight.

For sulAp-mCherry strains, an overnight culture was diluted in complex medium and grown to at least OD_600nm_ = 0.05 at 30°C before 0.1% arabinose was added. Samples were taken after 2, 4, 6 and 24 hours. After the two hours incubation, a subculture was treated with AT (10 µg/mL) and samples were taken after 2, 4 and 22 hours of growth at 30°C. Samples were re-suspended in PBS and Relative Fluorescence Units (RFU) of mCherry were detected with the FLUOstart micro-plate reader (BMG Labtech). The final RFU value was determined by normalising to cell density (OD_600nm_) then subtracting the normalised RFU of the untreated control from the same time point.

### 2-D DNA agarose gel electrophoresis and Southern hybridisation

Samples of cells were taken at the indicated time points and the DNA prepared as previously outlined (44, 70). DNA was digested with either EcoRI (array region) or HincII (region upstream of the array). 2-D gel conditions and Southern hybridisation were as described by (70). Statistical analysis was determined using the Student’s t test.

## Supporting Information

**Fig. S1. Replication fork collapse and reversal in a *dnaB*ts mutant upon shift to a non-permissive temperature.** Cells were grown at 30°C in the presence of 0.1% arabinose for 2 hours to block replication at the array and then shifted to 42°C to inactivate the replisome. (A) DNA structures within the array region were visualised by 2-D gels from samples taken at 15-minute intervals following the shift to 42°C. (B) Percentage of Y-structured DNA within the array region at the specified time points. n = 3 (*C*) 2-D gel analysis of replication intermediates detected in the region immediately upstream of the array.

**Fig S2. Replication fork blocks persist over 24 hours in WT cells, SOS is not induced in a *recA* mutant but is induced after 4 hours in an *uvrD* mutant.** A) Representative images of WT cells following arabinose induction over time. Green foci indicate blocked replication and red cells indicate SOS induction. Scale bar = 1 micron B) mCherry fluorescence observed in *uvrD*- over time in blocked cells C) mCherry fluorescence emission at indicated time points in Relative Fluorescence Units (RFUs) in blocked and released cells for both *recA* and *uvrD* mutants. Scale bar = 4 micron. Biological replicates = 3.

**Fig. S3. *recG* mutant impairs replication restart.** Mutant strains underwent treatment as described for strains shown in Figure 6 but with cells containing the wild type DnaB helicase (i.e. not *dnaBts*). Replication intermediate structures visualised by 2-D gel analysis in the array and upstream regions (*A*), cell viability (B) and foci percentages (*C*) under indicated conditions. Note the Δ*recG* strain has higher levels of HJ remaining 10 minutes after the addition of AT than other strains. n = 2–3

**Fig. S4. Cell viability is restored in *dnaB*ts mutant combinations following replication restart.** Viability was obtained under the indicated conditions as in Figure 6. –ara, cells with no replication blockage. +ara +AT 30°C, cells with replication blocked for 2 hours and then then released. + ara + AT TS, cells blocked for 2 hours, shifted to 42°C for 1 hour and then returned to 30°C and AT added. n = 2–3

**Table S1.** *E. coli* strains used in this study

